# Peripheral ghrelin administration prevents the behavioral effects of restraint stress in mice: possible implication of PVN^CRH^ neurons

**DOI:** 10.1101/2022.05.26.493640

**Authors:** Raoni Conceição Dos-Santos, Rafael Appel Flores, Aline Alves de Jesus, Rodrigo Rorato, André Souza Mecawi, José Antunes-Rodrigues, Lucila Leico Kagohara Elias

## Abstract

Ghrelin is a gut-derived hormone that is secreted during conditions of negative caloric balance and acts as a key modulator of feeding, increasing food intake and affecting several physiological systems such as metabolism, behavior and the control of endocrine and autonomic functions. Previous studies showed that ghrelin participates in the stress response, acting on hypothalamic paraventricular nucleus neurons that express corticotropin-releasing hormone (PVN^CRH^ neurons). In the present study, we investigated the effects of ghrelin administration on the behavioral responses to restraint stress in mice. In their homecage, C57Bl6 mice in basal conditions expressed the behaviors of surveying, walking, rearing, grooming and, to a lesser extent, digging, climbing and freezing. Restraint stress increased the time spent in grooming without significant changes in other behaviors. Ghrelin administration did not affect behavior in control mice, but it reversed the effect of restraint stress on grooming. Chemogenetic activation of PVN^CRH^ neurons by clozapine N-Oxide (CNO) administration in hM3Dq DREADD mice increased grooming, while ghrelin mitigated this effect. In addition, CNO administration decreased walking and rearing, both in the presence or absence of ghrelin. Food intake was increased by ghrelin administration, however, it was not affected by stress or CNO. These results indicate that ghrelin decreases the activity of PVN^CRH^ neurons, partially preventing the behavioral effects of restraint stress. The inhibitory input to PVN^CRH^ neurons probably arrives from other nuclei, since GABAergic neurons were not identified in the PVN neurons of these mice.

## INTRODUCTION

The brain monitors energy balance, changing feeding behavior and energy expenditure as necessary to maintain metabolic status. In response to changes in neural and hormonal cues related to caloric deficit animals feel hunger, an unpleasant sensation that motivates food intake^1^. In an ethological context, feeding behavior may be observed in three parts: the experiencing of hunger; the search for food; and the consumption of food^2^. After exposure to hunger-inducing stimuli, such as food deprivation or food restraint, rodents often increase the exploration of the environment despite the exposure to potential harm, a behavioral response often associated with a decrease in anxiety-like behavior^3–6^. In this context, ghrelin is a gut-derived hormone that is secreted during conditions of negative caloric balance and acts as a key modulator of feeding, increasing food intake and affecting several physiological systems such as metabolism, behavior and the control of endocrine and autonomic factors^7^.

The effects of intracerebroventricular (ICV) administration of ghrelin on the plasma hormone concentrations^8,9^ and autonomic regulation^10,11^ suggest that the hypothalamic paraventricular nucleus (PVN) participates in the responses elicited by ghrelin. We have previously demonstrated that ghrelin changes the activity of PVN neurons, exciting or inhibiting different subpopulations of neurons, and that the inhibitory effects are not expressed in the presence of γ-aminobutyric acid (GABA) and glutamate receptor antagonists^12^. These results suggest that the direct effects of ghrelin are excitatory and that the inhibitory effects are dependent on the activation of an inhibitory circuit.

The PVN is pivotal for the endocrine control of the stress response. In response to stress, neurons in the PVN release corticotropin releasing hormone (CRH) in the median eminence, this hormone reaches the anterior pituitary and induces the secretion of adrenocorticotropic hormone in the blood, which in turn stimulates the release of glucocorticoids from the adrenal gland^13^. ICV administration of CRH inhibits food intake^14,15^ and we have shown that ghrelin inhibits PVN^CRH^ neurons^12^, thus decreased activity of anorexigenic PVN^CRH^ neurons might contribute to the orexigenic effects of ghrelin. Additionally, previous studies showed that ghrelin participates in the stress response, likely through an effect on PVN^CRH^ neurons^16^.

PVN^CRH^ neurons also participate in immediate responses to stress that occur prior to changes in the plasma levels of ACTH^17^. This observation indicates that, in addition to the control of the hypothalamus-pituitary-adrenal axis, subpopulation of PVN^CRH^ neurons are involved in other stress-related responses. Corroborating this hypothesis, PVN^CRH^ neurons form synapses with other PVN neurons, influencing additional responses to stress^18^ and project to diverse brain regions such as the *globus pallidus*, involved in the control of locomotion^19^ and the pre-sympathetic autonomic neurons that control the autonomic tonus^20^. Previous studies have postulated that neuroendocrine and non-neuroendocrine PVN^CRH^ neurons may be activated independently^20,21^. Taken together, these observations suggest that, in addition to the well-known endocrine effects, PVN^CRH^ neurons may directly participate in several physiological responses and that different subpopulation of these neurons may be independently controlled.

Food deprivation changes the hierarchy of behavioral control, so that hungry animals prioritize food seeking in lieu of remaining in a safe environment, however, few studies have investigated the impact of concurrent competing stimuli in the ethological outcome^2^. It is plausible that ghrelin triggers a behavioral shift from passive stress coping mechanisms, such as grooming and freezing, to increased environmental exploration related to food seeking behavior. To test this hypothesis, in the present study we investigated the effects of ghrelin administration and/or restraint stress in the homecage behavior of mice. Next, we assessed the effects of peripheral ghrelin administration on FOS expression in PVN^CRH^ neurons. Lastly, we used chemogenetics to test the effects of the activation of PVN^CRH^ neurons in the homecage behavior and the influence of ghrelin administration on these effects.

## MATERIALS AND METHODS

### Animals

All procedures were approved by Ribeirão Preto Medical School – University of São Paulo (FMRP-USP) ethics committee (200/2019) and are in accordance with current legislation. Mice used in this study were acquired from the central animal facility (C57Bl6) or bred in-house CRH-CRE (B6(Cg)-Crhtm1(cre)Zjh/J; Jackson Laboratories) and Rosa26 (B6;129-Gt(ROSA)26Sortm2Sho/J; Jackson Laboratories) mice. All animals were maintained in collective cages (up to 4 animals each), 12/12 hrs light/dark cycle (lights on between 06:00 and 18:00), room temperature of 22 ± 2 ºC and *ad libitum* access to water and food. All experiments were performed in male mice. In order to minimize the effects of unspecific stress, all animals were handled for three days prior to experimentation.

### Breeding of transgenic mice

To investigate the participation of CRH neurons in the regulation of food intake and stress-related behaviors CRH-CRE^+/+^ mice were used. To generate these animals two breeders of CRH-CRE^+/+^ (B6(Cg)-Crhtm1(cre)Zjh/J, 012704, Jackson Laboratories) were formed. The homozygosis for CRH-CRE was confirmed through genotyping with PCR. Then, the offspring from these two couples was used to generate new pairs, so that there was no mating between offspring from the same parents, in order to avoid the interference of familial effects. The offspring from these breeders was used in the chemogenetics experiments.

Neuroanatomical analysis of CRH neurons is seldom possible through traditional techniques of immunofluorescence and immunohistochemistry, thus we generated transgenic mice that expressed enhanced green fluorescent protein (EGFP) in CRH neurons. For this purpose, CRH-CRE^+/+^ mice were bred to Rosa26-EGFP^+/+^ (B6;129-Gt(ROSA)26Sortm2Sho/J, 004077, Jackson Laboratories) mice, the offspring was then used for the immunofluorescence experiments.

### Analysis of the behavioral changes induced by ghrelin administration or restraint stress

To habituate the mice (n = 37, 8-11/groups) to the homecage, they were maintained in individual cages for 3 days prior to the behavioral experiments. During this period, the same researcher handled these mice and moved their cages individually to the recording room, in order to habituate these mice to the experimental conditions and decrease the influence of unspecific stress. On the day of the experiment, food and water were briefly removed from the cages and the mice received either vehicle (sterile 0.9 % NaCl, i.p.) or ghrelin (0.4 mg/kg; Bachem, USA; diluted in 0.9 % NaCl, i.p.) administration. Immediately after the injection, the animals were either returned to their homecage for 30 minutes or subjected to 30 minutes of restraint stress prior to returning to their homecage. After this period, the homecage was moved to an adjacent room and the animal behavior was recorded for 15 minutes. A representative schematic illustration of the experiment is provided (**Fig. 1A**). After recording (45 minutes after injection), the homecages were moved back to the ventilated rack and a pre-weighed pellet of chow and the water bottle were made available to the mice. Two hour later, the chow pellet was weighed to measure food intake.

**Figure 1.**
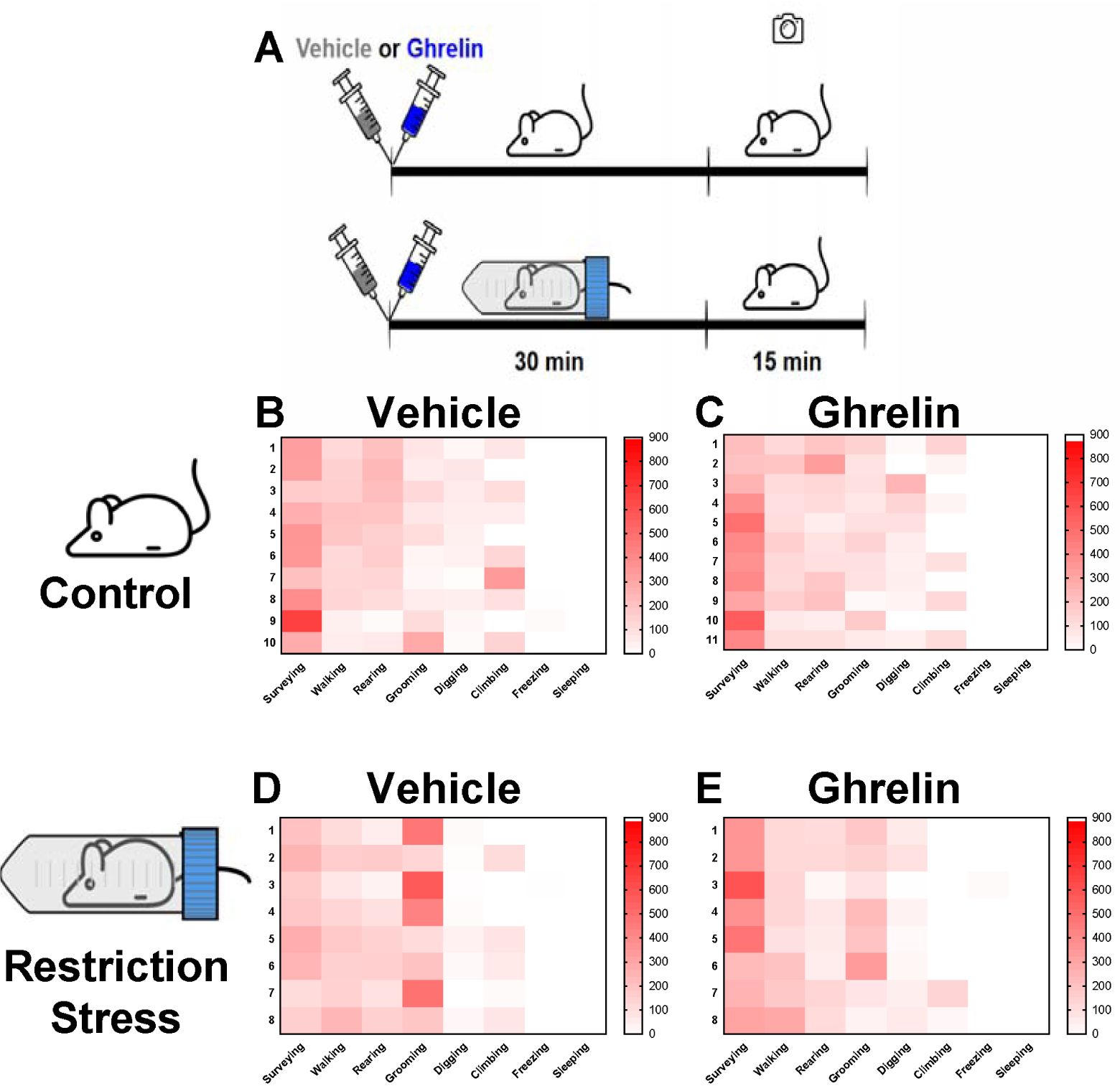
Schematic timeline of the experiment (**A**). Qualitative analysis of a heat-plot of expressed behaviors showed that control mice exhibited mostly exploratory activities (surveying, walking and rearing), and lesser time was spent grooming, digging or climbing (**B**). Restraint stress (**D**) increased time spent in grooming, mostly by decreasing surveying. Ghrelin did not change the behavioral pattern in control mice (**C**), but decreased the expression of grooming after stress (**E**). Freezing and sleeping were not observed in this experiment.

The ghrelin dose used in this experiment was determined by previous studies that demonstrated that this is the minimal dosage that changes food intake and anxiety-like behavior in mice^22,23^. All experiments were carried out in the morning, between 07:00 and 11:00, a period in which the mice generally do not eat significant amounts of food^24^, so that the orexigenic effects of ghrelin could be observed.

### Restraint stress

The restraint stress was carried out as in previous studies^25^, mice were placed in a manufactured restraint apparatus. The apparatuses were made from polypropylene 50 mL Falcon tubes, holes were drilled in the side of the tubes and the tip was cut, allowing ample ventilation to the animal. A hole was also drilled in the lid of the tube and the mouse’s tail was fitted through this hole, contributing to the restraint of the mice by hindering it from turning inside the tube. The mice were placed in the tube, which was positioned in a small support to avoid rolling. The animals were kept in this condition for 30 minutes. Restraint stress was used since it does not cause pain or physical harm to the animals, being classified as a psychophysical stress model^25^. Although restraint stress is usually maintained for long periods (between 1 and 6 hours), 30 minutes are sufficient to change hypothalamic levels of mRNA^26^ and change physiological functions, such as stress-induced hyperthermia^27^. Therefore, the model used causes the least possible discomfort to the animal, constituting a mild psychophysical stressful stimulus.

### Behavioral analysis

The videos were analyzed by an investigator blinded to the experimental groups, following previously described protocols^17^, with minor alterations. Füzesi et al.^17^ demonstrated that mice perform eight behaviors when observed in their homecage: surveying, walking, rearing, grooming, freezing, digging, sleeping and chewing. In that study, the researchers replaced the grid that covers the cage with an acrylic lid and did not remove the chow. In our study, we chose to keep the original grid, in order to decrease the effects of novelty stress on the behavioral response, consequently, the mice could climb the grid. In addition, we removed the food, so that chewing behavior could not be observed.

To quantify behavioral parameters, the time spent in each activity was registered. ‘Walking’ was scored when the mice moved horizontally through the cage. ‘Rearing’ was marked when the mice dislocated the weight to the hindpaws and stretched the anterior portion of the body vertically. ‘Grooming’ was considered when the animals repetitively licked the paws and back in stereotypical fashion. When the animal was not walking or rearing, but interacting with the environment (sniffing, listening or looking to the ambient, etc.) the behavior was considered as ‘surveying’. When the animal was completely immobile the behavior was measured as ‘freezing’. When the mice moved the bedding to explore the bottom of the cage, we considered the behavior as ‘digging’. When the mice climbed to the cage grid the behavior was considered as ‘climbing’. None of the animals slept during the recording period in this study. A representative video of the behaviors observed is provided (**Supplementary Movie 1**).

### Effects of ghrelin on FOS expression in the PVN CRH and GAD67 neurons

CRH-EGFP mice (n = 4, 2/group) were used for the qualitative analysis of FOS expression in CRH or GAD67 neurons in the PVN after ghrelin administration. These mice underwent administration of vehicle or ghrelin. Two hours after injection, mice were anesthetized with an overdose of ketamine (Syntec, BR) + xylazine (Syntec, BR) mixture (respectively, 200 mg/kg and 20 mg/kg), then the chest cavity was opened and the heart exposed, a needle was inserted in the cardiac apex and the right atrium was cut. Then, the animals were perfused with saline solution (0.9 % NaCl) for one minute, followed by 10 % formalin for five minutes. Then, the skulls were removed and the brain was obtained and post-fixed in 10 % formalin for 4 hours. After which, the brains were stored in 30 % sucrose (4 °C). Lastly, the brains were cut in 30 µm slices in a cryostat (Micron HM525E) and stored in anti-freezing solution for posterior processing.

### Immunofluorescence

For the immunostaining of GAD67 and FOS, the brain slices were washed three times for 10 minutes (0.01 M PBS), then incubated in blocking solution (5 % normal horse serum and 0.3 % Triton, diluted in 0.1 M PBS) for 60 minutes. After which, the slices were incubated overnight in a mixture of primary antibodies anti-FOS (rabbit anti-FOS, 1:10000, cat: p38, Calbiochem, USA) and anti-GAD67 (mouse anti-GAD67, 1:1000, cat. MAB5406, Sigma-Aldrich, USA) diluted in blocking solution. On the following day, the slices were washed three times for 10 minutes (0.01 M PBS), then incubated with secondary antibodies (goat anti-rabbit conjugated with Alexa 594, 1:500, cat. A11037, Invitrogen, USA; and donkey anti-mouse conjugated with Alexa 488, 1:500, cat. 715545151, Jackson Immunoresearch, USA). Next, the slices were washed three times for 10 minutes (0.01 M PBS) and mounted with Fluoromount (Thermofisher, USA). Images were collected in a Leica microscope.

CRH-EGFP mice were used for endogenous fluorescence of CRH neurons, and FOS was stained as described above. In CRH-CRE^+/+^ mice with DREADD-G_q_ expression, FOS staining was carried out as above, with the substitution of the primary antibody and secondary antibodies (respectively, goat anti-FOS, 1:1000, cat: AB87655, Abcam, USA; and donkey anti-goat conjugated with Alexa 488, 1:500, cat. A11055, Molecular Probes, USA), mCherry was endogenously expressed.

### Analysis of the behavioral changes induced by activation of PVN^CRH^ neurons or by the excitation of PVN^CRH^ neurons after ghrelin administration

CRH-CRE^+/+^ mice (n = 6) were microinjected with CRE-activated adeno-associated (AAV) viruses (pAAV-hSyn-DIO-hM3D(Gq)-mCherry; serotype 5; titter ≥ 3×1012 UNC Vector core, Chapel Hill, NC) to induce the expression of G_q_-associated DREADDs in the PVN (**Fig. 2A**). DREADDs (Designer Receptors Exclusively Activated by Designed Drugs) are modified receptors that are exclusively activated by selective drugs that do not affect endogenous receptors^28^.

**Figure 2.**
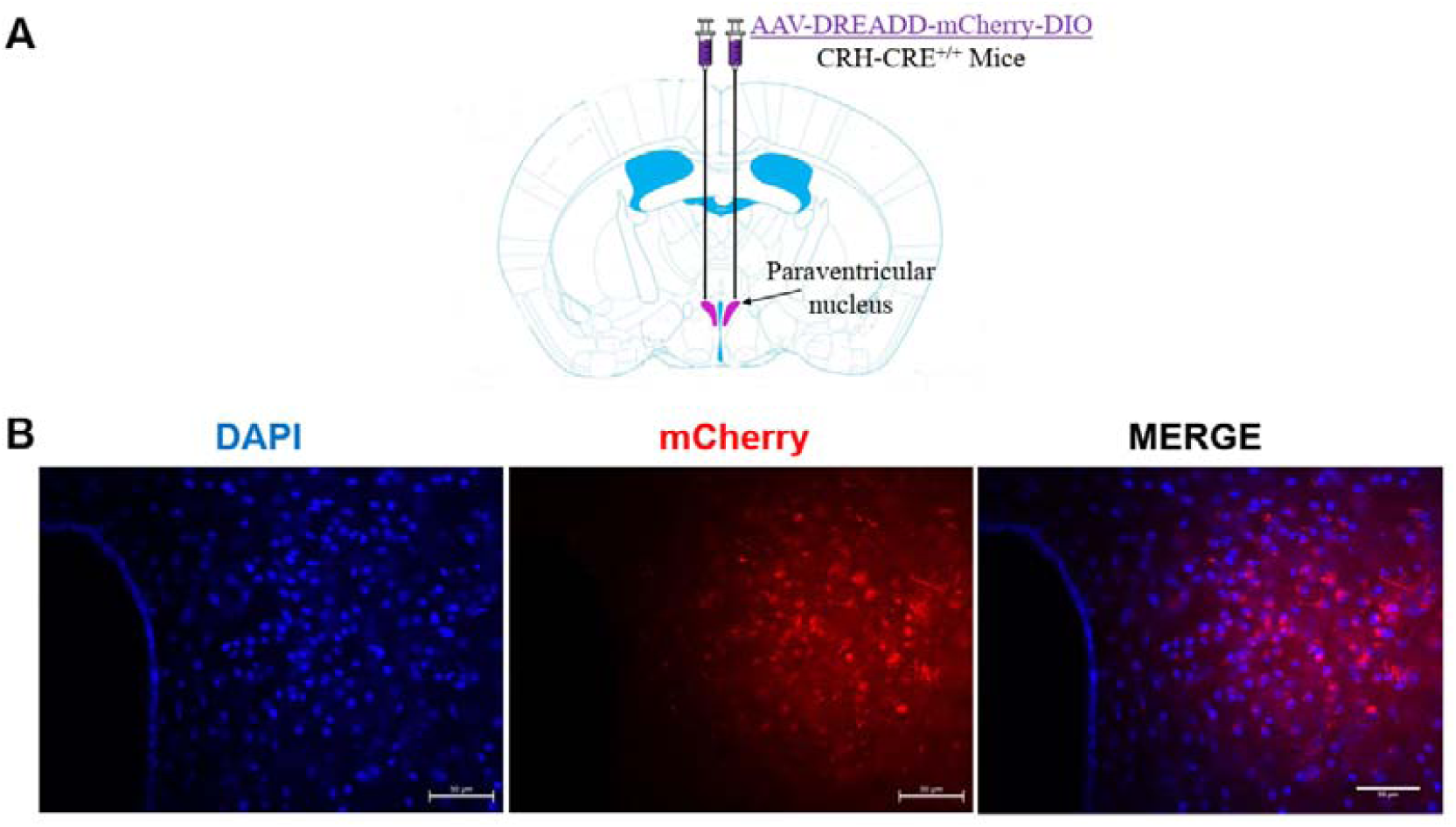
Schematic illustration of the microinjection of AAV in the PVN of CRH-CRE^+/+^ mice (**A**). Representative photomicrograph of DAPI and mCherry staining in the PVN of a CRH-CRE^+/+^ mice (**B**).

To perform the microinjection, borosilicate glass micropipettes were manufactured in a PIP6 vertical pipette puller (HEKA Electronics Inc, USA). The micropipette was placed in a stereotaxic device and used for the administration of AAVs. For the stereotaxic surgery, the mice were anesthetized with a mixture of ketamine and xylazine (respectively, 100 mg/kg and 10 mg/kg). Then, the animals were placed in a stereotaxic frame with digital displays (Kopf Instruments, USA). The coordinates used for the PVN were: 1.4 mm posterior to the bregma; 0.25 mm lateral to the midline and 4.9 mm deep in relation to the skull^29^. A small hole was drilled in the skull along the midline and the micropipette was lowered to the specified coordinates, then 60 nL of the AAV were slowly delivered to the nucleus. After administration we waited for 5 minutes prior to the removal of the micropipette to allow the diffusion of the AAV. Then the same procedure was repeated on the other side for bilateral microinjection. After viral vector delivery, the hole in the skull was filled with bonewax (Medline, BR), the skin was sutured and mice received a prophylactic dose of flunixin meglumine (1 mg/kg, s.c.; MSD Saúde Animal, BR)..

Four weeks after AAV microinjection, the behavioral experiments were carried out; the same habituation procedure described above was performed. On the day of the experiment, food and water were removed, then, the animals were handled and replaced in their homecage and allowed to rest for 30 minutes. Next, they were moved to the adjacent recording room and baseline behavior was recorded for 15 minutes. After which, the cages were transferred back to the ventilated racks and food intake was measured for two hours. On the following day, mice received an injection of Clozapine N-oxide (CNO; Tocris, USA, 1 mg/kg, i.p.), 30 minutes after drug administration they underwent the same procedures. Lastly, 7 days after the previous experiments, the mice received a second injection of CNO, immediately followed by an injection of ghrelin. 30 minutes after the administration of these drugs, they were submitted to the same behavioral procedures. Previous studies demonstrated that the dose of CNO used does not cause clozapine-like effects in mice^28^, indeed a 10x higher dose is necessary for CNO to induce off-target effects in mice and rats^30^

At least 4 days after the behavioral experiments, mice received another CNO injection. After two hours, the animals were anesthetized and submitted to transcardiac perfusion and the brains were removed and sectioned in the cryostat, as detailed above. The brain slices were mounted with Fluoromount with DAPI (Thermofisher, USA) and taken to a fluorescence microscope. Next, we performed FOS immunostaining to confirm the capacity of CNO to induce the activity of mCherry positive PVN neurons.

### Statistical Analysis

Graphpad Prism v. 7.0 software was used for all statistical analyses. The effects of ghrelin and restraint stress in C57Bl6 mice were compared by two-way ANOVA followed by Tukey’s post-test, when appropriate. One-way ANOVA for repeated measures followed by Tukey’s post-test was used to compare the differences between the behavior of DREADD-G_q_ mice. Additional behaviors from DREADD-G_q_ mice were not normally distributed, thus were compared with Friedman’s test, followed by Dunn’s post-hoc test. All data are described as means ± SEM.

## RESULTS

To identify the behavioral patterns of mice in their homecage, we performed a qualitative analysis of the time spent in each behavior through a heatplot (**Fig. 1B-E**). Control mice (**Fig. 1B**) spent most of the time surveying, walking and rearing, while less time was spent in the other behaviors, similarly to the results found in other studies^17^. Ghrelin administration (**Fig. 1C**) did not change the behavioral pattern observed when compared to the control-vehicle group. On the other hand, mice submitted to restraint stress (**Fig. 1D**) showed an increased time spent grooming at the expense of surveying. Interestingly, ghrelin pretreatment blunted the increase of grooming induced by restraint stress (**FIG. 1E**). Digging and climbing were not consistently performed by the mouse. Freezing was seldom expressed and none of the subjects slept during the recording.

The expression of mCherry in the PVN was used to confirm DREADD expression in PVN^CRH^ neurons (**Fig. 2B**). Next, we quantitatively assessed the influence of restraint stress and ghrelin on homecage behavior (**Fig. 3**), especially the behaviors directly affected by PVN^CRH^ neurons: walking, rearing, grooming and freezing^17^. This analysis revealed that walking (**Fig. 3A**) and rearing (**Fig. 3B**) were not significantly different between experimental groups. Time spent in grooming (**Fig. 3C**) was affected by both stress (F_(1,33)_ = 17.31; p < 0.001) and ghrelin treatment (F_(1,33)_ = 4.82; p = 0.035), with significant interaction between both factors (F_(1,33)_ = 5.59; p = 0.024). Post-hoc analysis demonstrated that grooming was significantly increased in the vehicle-stress group when compared to all other groups (p < 0.05). Therefore, psychophysical stress increases the time spent in grooming, and the administration of ghrelin may mitigate this effect. Food intake (**Fig. 3D**) was affected by ghrelin (F_(1,33)_ = 5.64; p = 0.023), as expected, with no differences caused by stress (F_(1,33)_ = 0.51; p = 0.48) or interaction between factors (F_(1,33)_ = 0.088; p = 0.77). Additional behavioral data is shown in **Table I**.

**Figure 3.**
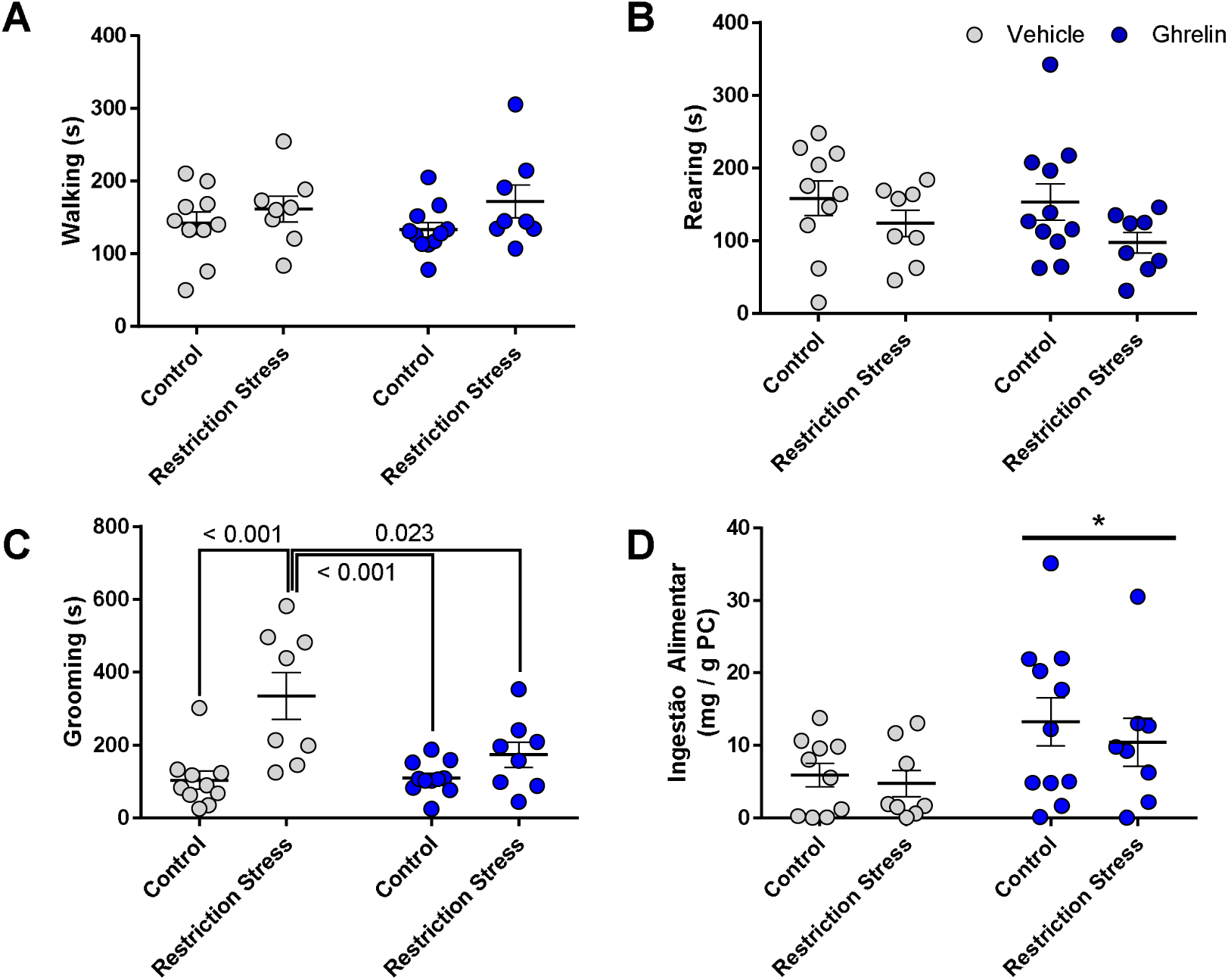
Walking (**A**) and rearing (**B**) behaviors were not affected by either ghrelin administration or restraint stress. Restraint stress increased time spent in grooming, an effect that was reversed by ghrelin administration (**C**). Ghrelin increased food intake, and restraint stress did not affect this response (**D**). * p < 0.05 vs vehicle.

**Table I.**
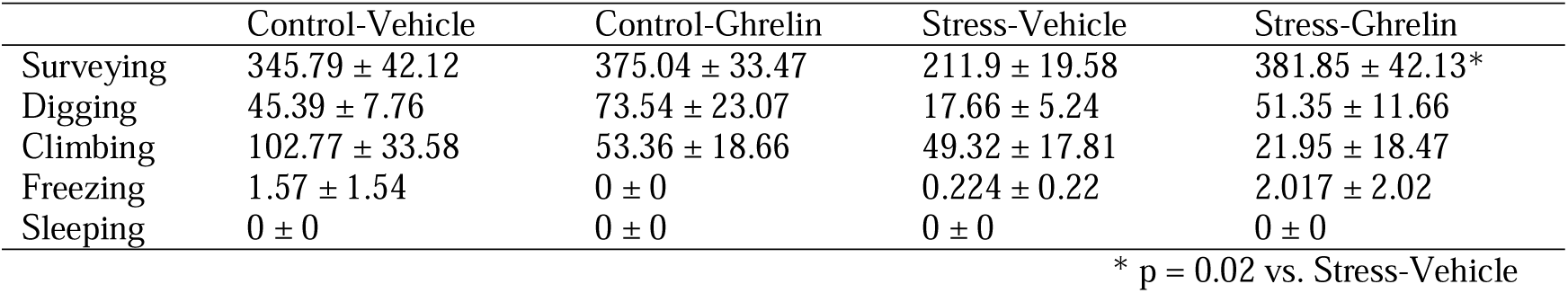
Additional behavioral data

|  | Control-Vehicle | Control-Ghrelin | Stress-Vehicle | Stress-Ghrelin |
| --- | --- | --- | --- | --- |
| Surveying | 345.79 ± 42.12 | 375.04 ± 33.47 | 211.9 ± 19.58 | 381.85 ± 42.13* |
| Digging | 45.39 ± 7.76 | 73.54 ± 23.07 | 17.66 ± 5.24 | 51.35 ± 11.66 |
| Climbing | 102.77 ± 33.58 | 53.36 ± 18.66 | 49.32 ± 17.81 | 21.95 ± 18.47 |
| Freezing | 1.57 ± 1.54 | 0 ± 0 | 0.224 ± 0.22 | 2.017 ± 2.02 |
| Sleeping | 0 ± 0 | 0 ± 0 | 0 ± 0 | 0 ± 0 |

\* p = 0.02 vs. Stress-Vehicle

Ghrelin administration does not induce FOS expression in CRH neurons in the PVN (**Fig. 4**), which indicates that these neurons are not activated by this treatment. Then, we investigated the expression of the 67 KDa isoform of the glutamic acid decarboxylase enzyme (GAD67). Neither GAD67 nor FOS expression were detected in the PVN neurons after intraperitoneal injection of vehicle or ghrelin (**Supplementary Fig. 1**).

**Figure 4.**
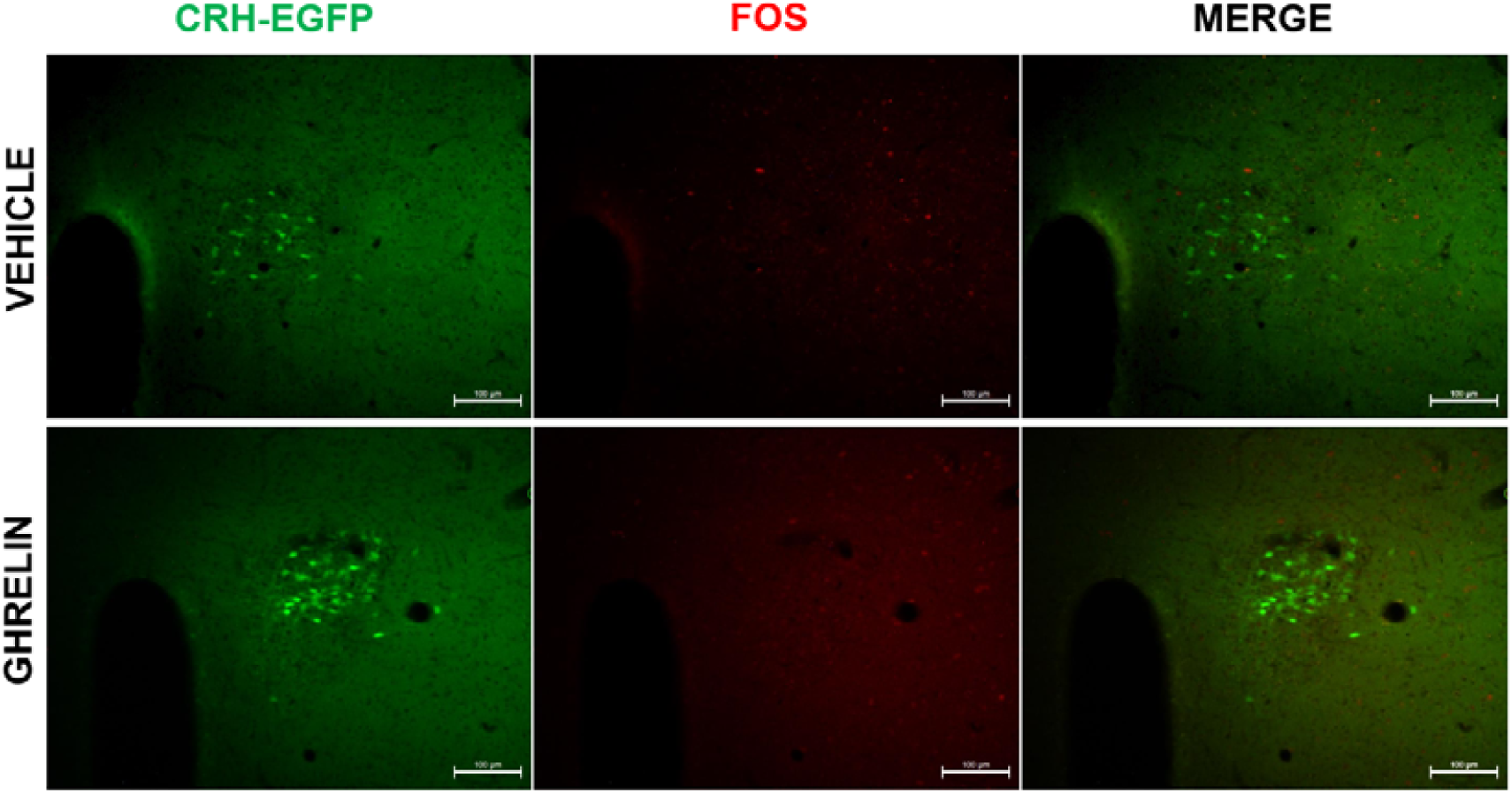
CRH-EGFP (green) neurons in the PVN did not express FOS (red) after vehicle or ghrelin administration in mice.

In CRH-CRE^+/+^ mice with the expression of DREADD-G_q_ in PVN^CRH^ neurons, the administration of CNO changed the animal behavior in the homecage (**Fig. 5**). A representative video of the behaviors observed is provided (**Supplementary Movie 2**). CNO significantly decreased the time spent walking (p < 0.001, **Fig. 5B**) or rearing (p = 0.001, **Fig. 5C**) when compared to baseline behavior. After the concomitant administration of CNO and ghrelin, the time spent walking (p = 0.003) and rearing (p = 0.021) are also significantly different from baseline. Time spent in grooming (**Fig. 5D**) was significantly increased after CNO administration in comparison to the baseline (p = 0.016). After concomitant administration of CNO and ghrelin, the time spent grooming was reversed, no longer being different from baseline. Thus, similar to the results demonstrated after restraint stress, ghrelin administration reversed the increase in grooming behavior after the chemogenetic activation of PVN^CRH^ neurons. In addition, comparison between time in grooming for vehicle-injected C57Bl6 mice vs. CNO-injected DREADD G_q_ CRH-CRE^+/+^ mice was significantly different (103.09 ± 24.7 vs. 217.2 ± 50.5, p = 0.039, Unpaired Student T-test), which indicates that the increase in grooming time is not due to the injection stress. Although the differences were not statistically significant, some animals demonstrated increased time spent in freezing behavior both after CNO or CNO+ghrelin administration (**Fig. 4E**). Food intake was not changed by any of the treatments investigated (**Fig. 4F**). Additional behavioral data is described in **Table II**. FOS expression in mCherry positive neurons within the PVN after CNO administration (**Supplementary Fig. 2**) confirms that CNO activated DREADDed neurons, as expected^28,31^.

**Figure 5.**
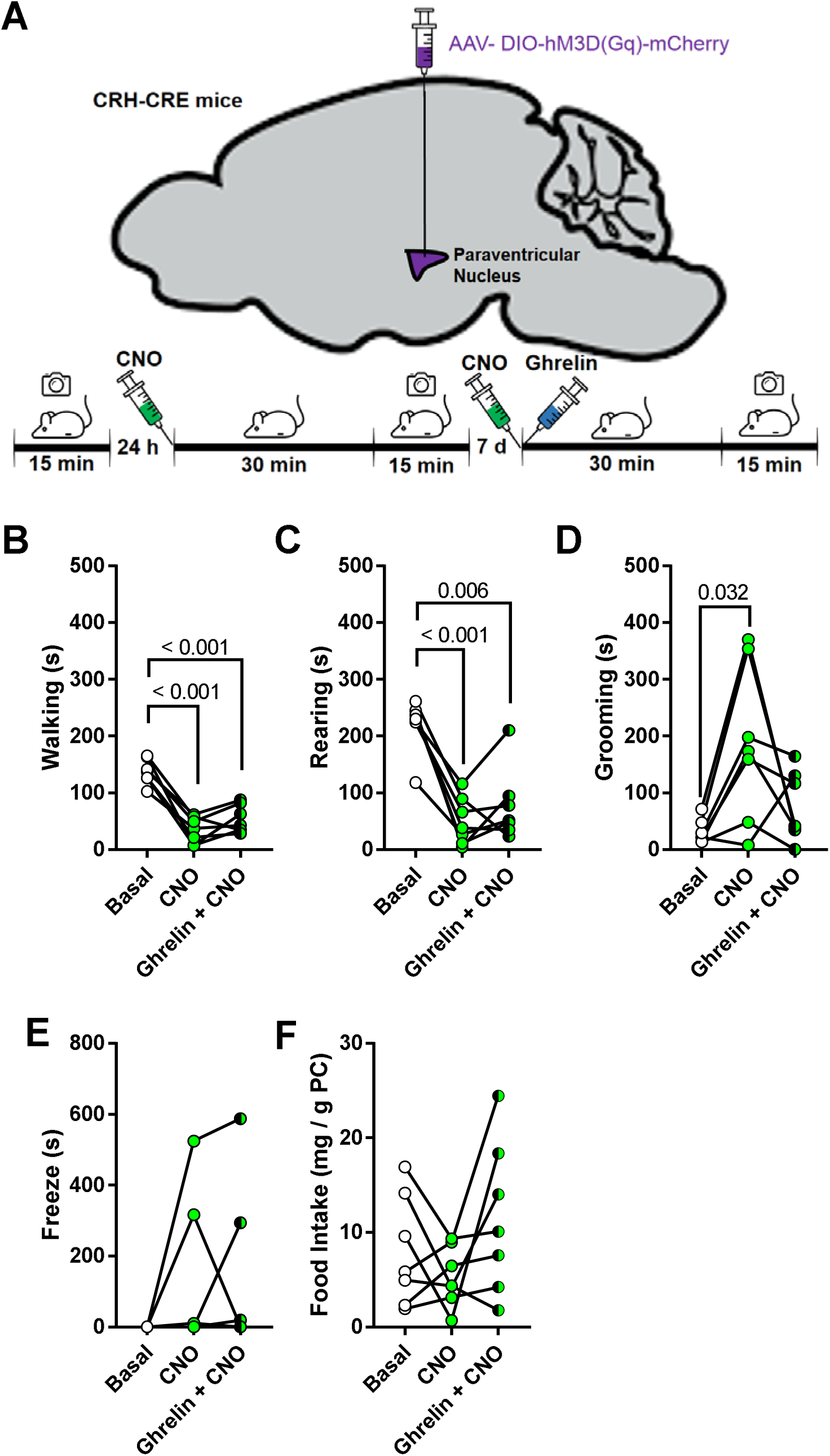
Schematic illustration of the microinjection of AAVs in the PVN of CRH-CRE mice and timeline of the experiment (**A**). Walking (**B**) and rearing (**C**) decreased after CNO and CNO + ghrelin administration. Grooming (**D**) increased after CNO, and this effect was reversed by CNO + ghrelin. Freezing (**E**) was not significantly affected, however, some animals demonstrated freezing behavior after CNO or CNO+ghrelin administration. Food intake (**F**) was not affected by the treatments.

**Table II.** Additional behavioral data from DREADD-G_q_ mice

|  | Basal | CNO | CNO + Ghrelin |
| --- | --- | --- | --- |
| Surveying | 410.9 ± 29.59 | 476 ± 60.62 | 515.5 ± 83.73 |
| Digging | 20.38 ± 8.82 | 16.68 ± 5.31 | 8.82 ± 4.28 |
| Climbing | 74.43 ± 22.99 | 3.96 ± 3.96 | 33.67 ± 33.67 |
| Sleeping | 0 ± 0 | 0 ± 0 | 0 ± 0 |

## DISCUSSION

In the present study we demonstrate that ghrelin modulates behavioral changes induced by restraint stress and this effect seems to be mediated by PVN^CRH^ neurons. In C57Bl6 mice, ghrelin increased food intake, as expected. However, restraint stress did not change food intake, contrary to other studies that showed that restraint stress decreases food intake in rodents^32,33^. These differences between studies may be due to differences in the time spent under restraint stress, and, consequently, in the intensity of the stressful stimulus. Possibly, the time of restraint stress used in this study was not sufficient to change food intake. Similarly, another study in mice showed that one hour of restraint stress does not change food intake, but that repeated exposure to one hour of restraint stress leads to decreased food intake^34^. In addition, chemogenetic activation of PVN^CRH^ neurons also did not change food intake. All experiments described were carried out in the morning period in which mice generally eat small amounts of food^24^, thus the low motivational drive to eat might have influenced these results. Possibly, an anorexigenic effect of stress or PVN^CRH^ neuron activation would be demonstrated if the experiments were carried out in the dark period.

In addition to the effects of ghrelin on food intake, this hormone influences the response to stress^35^. Ghrelin effects on the control of the behavioral response to stress are not yet completely understood, however, some studies indicate that ghrelin mitigates some effects of stress. For instance, ghrelin administration attenuates the increase in anxiety-like and depression-like behaviors in mice that underwent social defeat stress^36,37^. Furthermore, mice in which the ghrelin receptor gene was knocked out show an increase in these behaviors when compared to control mice^37,38^. However, the tests commonly used to investigate the behavioral responses, such as elevated plus maze, open field, light-dark box, etc., are themselves stressful stimuli that increase plasma corticosterone^39^. Thus, recent studies highlight the need to investigate behavioral responses in situations that more closely resemble a familiar environment for the mice^17,40^ and the mechanisms through which the outcome behavior is selected in the presence of concurrent stimuli^2^. In the present study, we investigated the effects of ghrelin on the behavior of mice in their original cage, in this context, ghrelin did not affect exploratory behaviors, grooming or freezing, however, ghrelin attenuated the effects of mild intensity psychophysical stress.

Diverse models of stress differently influence the behavioral response. In addition, the intensity of restraint stress varies due to the duration of the restraint. Previous studies showed that 15 minutes of restraint stress did not affect the exploratory behavior of mice in the open field test, however, 50 minutes of restraint caused an increase in exploratory activities that could be reversed by pharmacological blockade of corticosterone synthesis^25^. Similarly, footshock stress increased homecage exploratory activity in mice, in addition to increasing time spent in grooming, however, the expression of grooming after footshock was not affected by blockade of corticosterone synthesis^17^. Therefore, the increase in grooming after footshock stress seems to be independent of changes in corticosterone levels. In the present study, 30 minutes of restraint stress did not change exploratory behavior, however, significantly increased the time spent in grooming.

Several studies demonstrated FOS expression in the PVN of rats after peripheral administration of ghrelin, however, the co-localization with CRH was not investigated^41–43^. Previous studies demonstrated that peripheral ghrelin administration induced FOS expression in the PVN and increased plasma corticosterone in mice^44^, which indicates an excitatory effect of ghrelin on hypophysiotropic PVN^CRH^ neurons. However, we^12^ and others^45^ have demonstrated in *in vitro* studies that the effects of ghrelin on CRH neurons are inhibitory. In the present study, peripheral ghrelin did not increase FOS expression in the PVN^CRH^ neurons. Thus, an excitatory effect of ghrelin on CRH neurons is unlikely. Possibly, the corticosterone releasing effects of ghrelin are due to direct effects on the pituitary, rather than on hypothalamic neurons, as shown previously in the literature^7^.

In order to verify if the inhibition of PVN^CRH^ neurons was due to GABAergic interneurons within the PVN we investigated the expression of GAD67 in this nucleus. GAD67 is an enzyme that converts glutamate to GABA^46^, therefore its expression indicates that the neuron is GABAergic. GAD67 was identified by imunnostaining in the PVN of rats subjected to prenatal programming^47,48^ or colchicine pretreatment^49^, other studies, however, showed that the PVN does not express GAD67-positive cell bodies, but rather neuron terminals^50^. Indeed, in-situ hybridization studies demonstrated that GAD67 and GAD65 mRNA are present in regions adjacent to the PVN rather than on the PVN itself^51^. Consistent with these data in the literature, in the present study, we did not find GAD67-positive neurons within the PVN. This indicates that the inhibitory input to the PVN probably arises from other nuclei. Several forebrain nuclei might be relevant sources of GABAergic input to the PVN, among them the arcuate nucleus^51^. This nucleus plays a key role in the regulation of feeding and neurons in this nucleus are sensitive to ghrelin^7^. Possibly, the peripheral administration of ghrelin increases the activity of inhibitory arcuate neurons to the PVN^CRH^ neurons decreasing their activity and, consequently, decreasing the behavioral impact of restraint stress. Further studies, however, are necessary to elucidate the putative Ghrelin → arcuate → PVN^CRH^ circuit involved in the effects of ghrelin on stress response.

The optogenetic photostimulation of PVN^CRH^ neurons also increases grooming behavior^17^, which indicates that the increased activity of PVN^CRH^ neurons participates in the increased grooming after stress. Self-grooming behavior is an important ethological resource that participates in several physiological function in mammals, such as social behavior, cleaning of the fur or wounds and preparation to sleep; however, in anxious/stressful situations grooming is evoked as a form of passive coping^52^. Additionally, the expression of grooming in aberrant situations is often used as an indicator in animal models of diseases that are related to complex, repetitive and stereotyped movements, such as Parkinson’s^53^. In basal situations, grooming is a low priority activity, since it is not related to the response to any challenging external stimulus, i.e. this behavior is often expressed when there is no motivation to perform other more pertinent activities^52^.

In this sense, the relationship between PVN^CRH^ neurons and the stimulation of grooming possibly indicates that a subpopulation of these neurons affects the behavioral control. Indeed, different populations of PVN neurons are directly related to the determination of behavior in mice. Previous studies have demonstrated that PVN^CRH^ neurons are related to fear, feeding and the response to the administration of ghrelin^54^. We have previously demonstrated that ghrelin, through a pre-synaptic effect, inhibits the activity of CRH neurons in acute brain slices^12^. In this sense, some acute stressful stimuli increase plasma ghrelin in rodents, such as psychological stress^55^ and maternal separation in newborn pups^56^, so that this hormone might contribute to the inhibition of the hypothalamus-pituitary-adrenal axis and the reestablishment of normal corticosterone levels after stress. Concomitantly, ghrelin might inhibit the expression of stress-related behaviors, contributing to the recovery of baseline behavior. In support to this hypothesis, mice knockout to ghrelin^36^ or ghrelin receptor^37,38^ show that in the absence of ghrelin signaling the animals are more susceptible to the anxiety-inducing effects of stress.

In this study, we induced the activation of PVN^CRH^ neurons through chemogenetic techniques. In DREADD-G_q_ mice, 30 minutes after CNO administration the animals showed a marked increase in grooming behavior and decreased exploratory activity. The decrease in exploration is considered a reliable indicator of the behavioral stress response, in its most extreme form, the animals express freezing, in which all locomotor and exploratory activities are abolished^39^. Some of the DREADD-G_q_ mice expressed freezing after CNO administration, however, this response was not consistently demonstrated in all animals studied. This behavioral pattern differs from the one demonstrated in C57Bl6 mice submitted to the restraint stress in this study. Possibly, the stress caused by CNO administration was more intense than the stress evoked by 30 minutes of restraint, thus causing a more pronounced behavioral response. Ghrelin administration prevents the effects of the activation of PVN^CRH^ neurons on grooming; nonetheless, the exploratory activities remain diminished. Therefore, the administration of ghrelin might decrease the impact of the stressful stimulus, supporting the coping process, but does not prevent all the behavioral effects of stress caused by the activation of PVN^CRH^ neurons. Taken together, the present results and the data from the literature reinforce the role of ghrelin as a stress-coping biomarker^57,58^.

In conclusion, the activation of PVN^CRH^ neurons inhibited exploratory activities, such as walking and rearing, and increased passive coping behaviors, such as grooming and freezing. These effects are similar to those evoked by several stress models. Ghrelin mitigates these behavioral effects, which indicates that this hormone might cause the inhibition of this subpopulation of PVN neurons. Similarly, ghrelin mitigates the effects of a mild intensity psychophysical stress, which further supports that this hormone might affect stress response. Although ghrelin may directly affect PVN neurons, its effects on PVN^CRH^ neurons are likely to arise from inhibitory inputs from other regions.

## ACKNOWLEDGEMENTS

The authors would like to thank Maria Valci dos Santos and Milene Mantovani Mata for their outstanding assistance and technical support.

## AUTHOR CONTRIBUTIONS

RCS, LKE and JAR designed the study. RCS, RAF, AAJ performed experiments. All authors participated in data analyses or acquisition, drafted/revised the manuscript, approved the final version, and agreed to be accountable for all the aspects of the work.

---

**Supplemental Figure 1.**
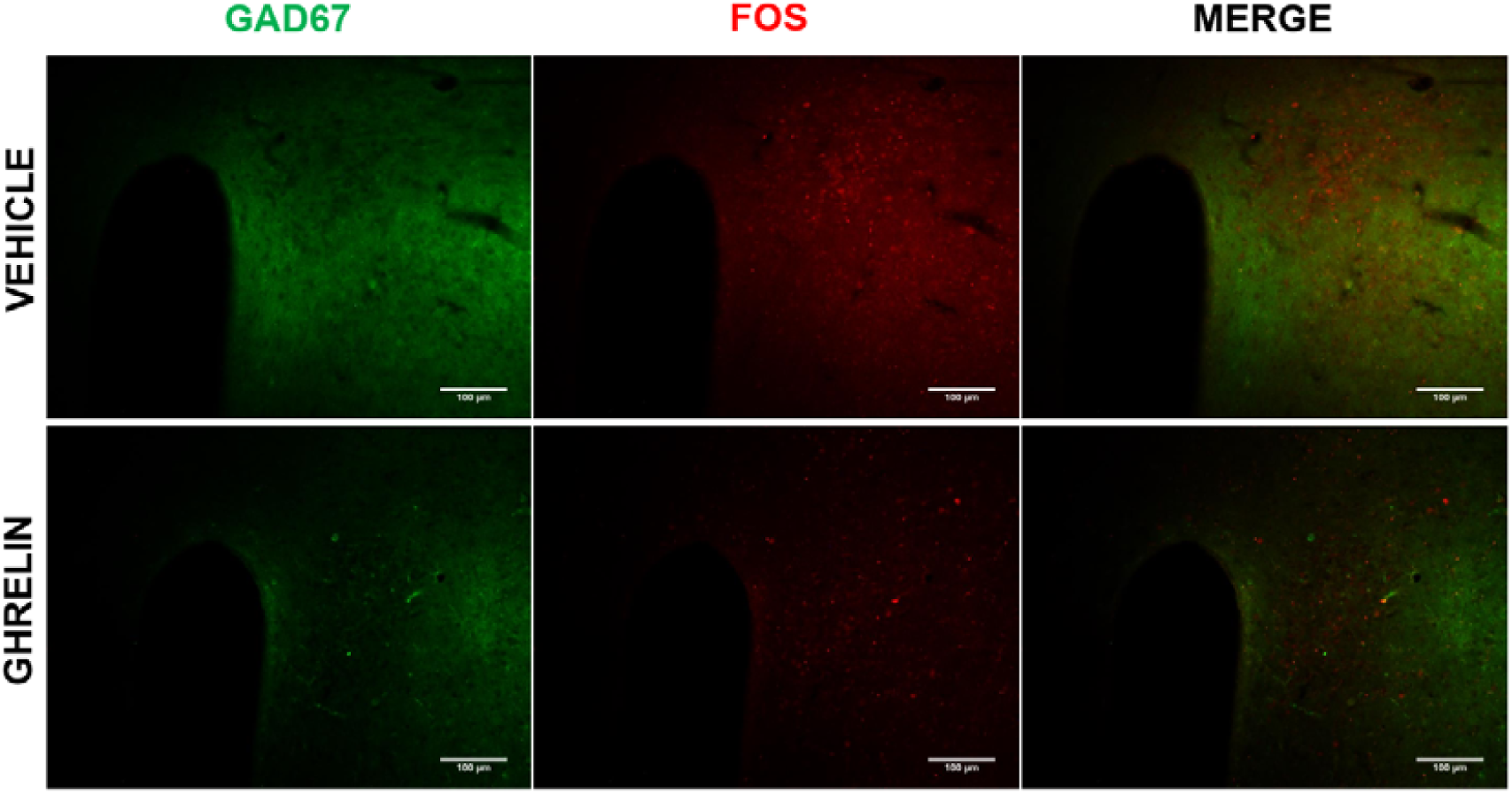
GAD67 (green) was not observed in the PVN. Few PVN neurons showed FOS expression (red) after vehicle or ghrelin administration.

**Supplemental Figure 2.**
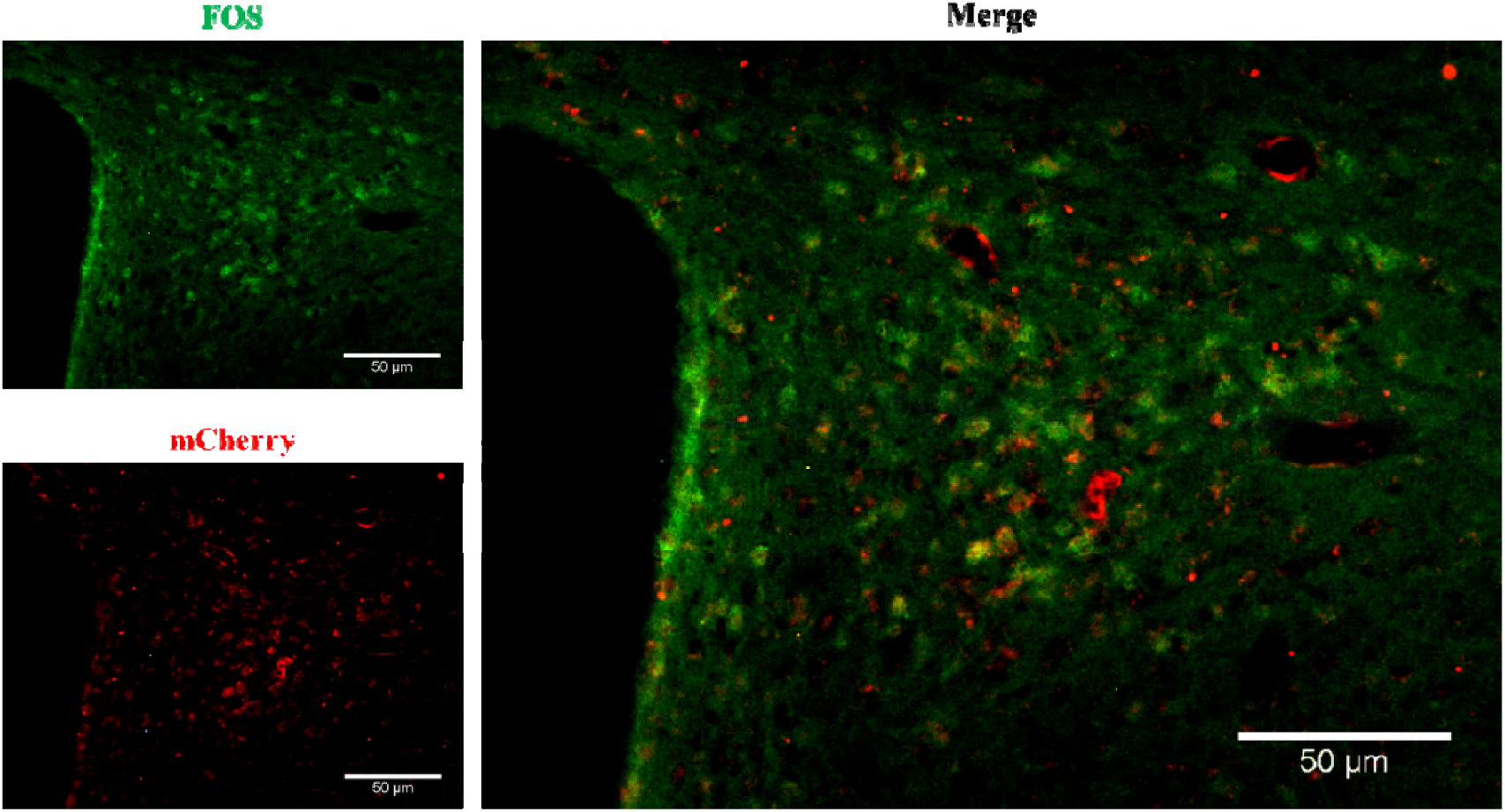
Representative figure of FOS expression (green) after CNO administration in CRH-CRE^+/+^ mice (mCherry, Red). Colocalization between FOS and mCherry (white arrows) highlights the activation of DREADDed PVN neurons after CNO administration.

