## Supplementary figures and images for "Peripheral ghrelin administration prevents the behavioral effects of restraint stress in mice: possible implication of PVN^CRH^ neurons"

### Supplementary Fig. 1

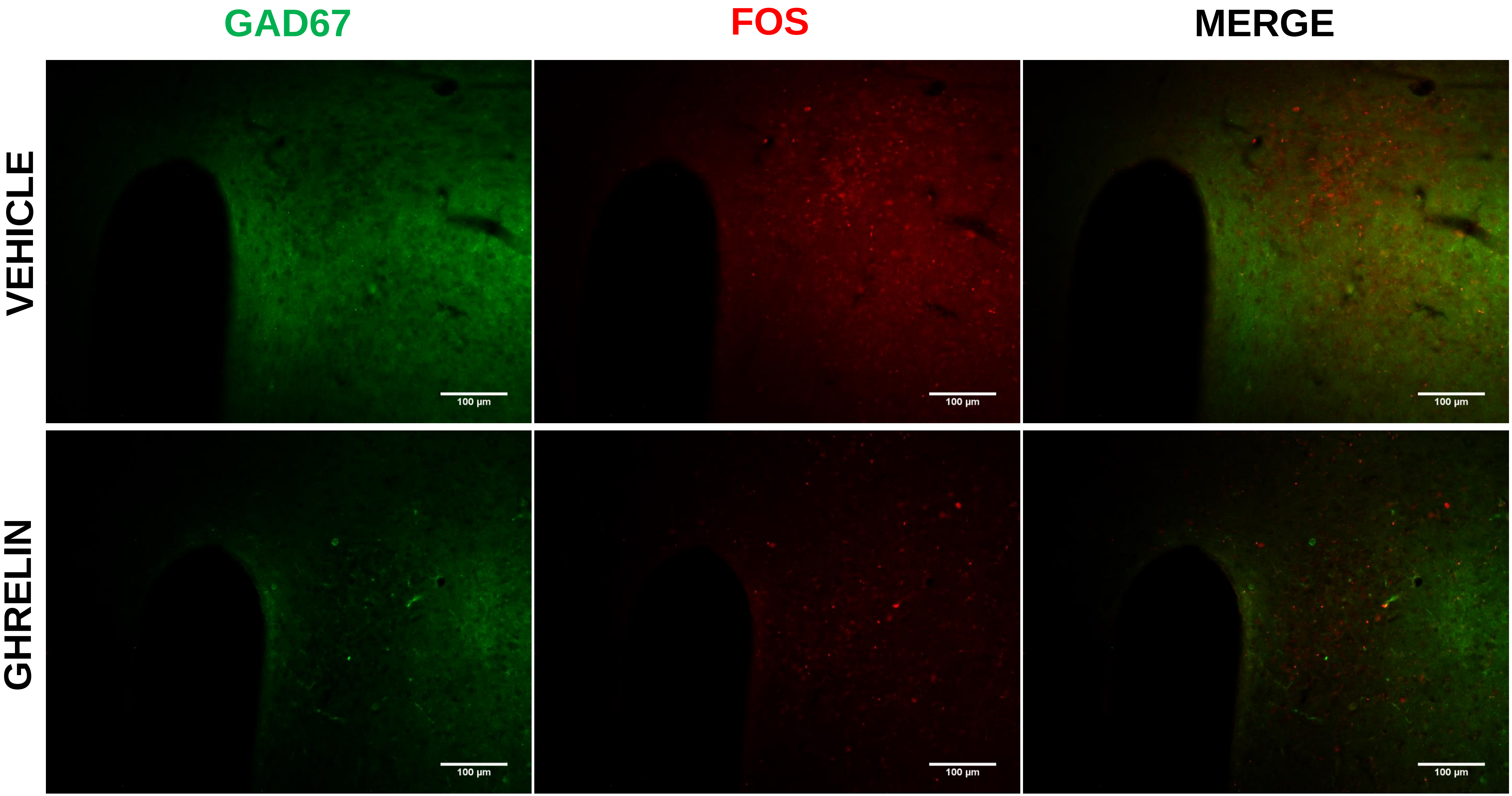

### Supplementary Fig. 2

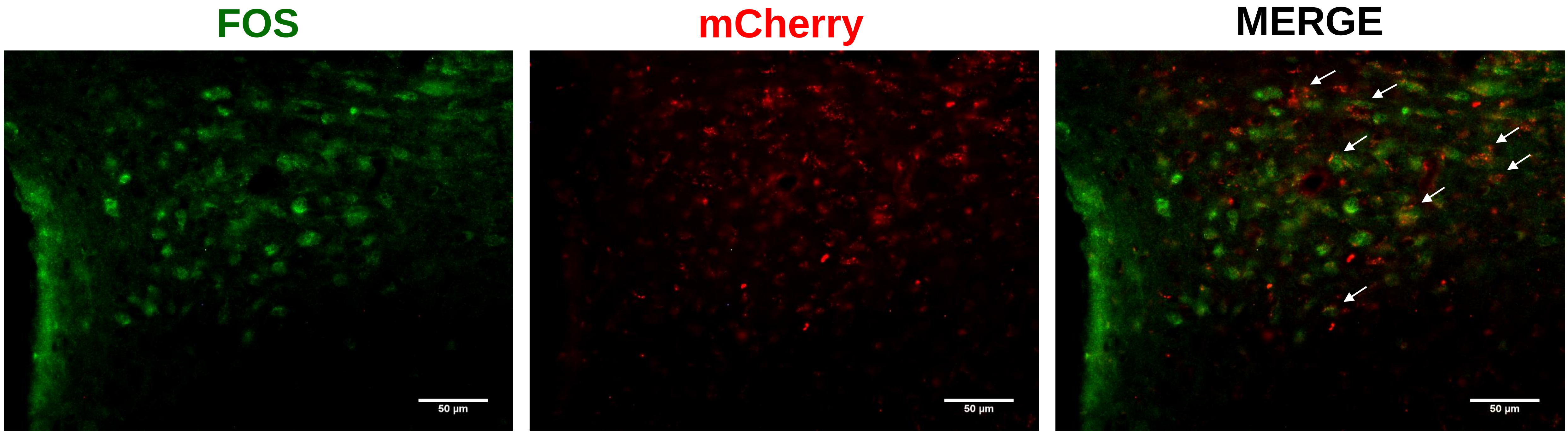
